# Evolution of a new testis-specific functional promotor within the highly conserved *Map2k7* gene of the mouse

**DOI:** 10.1101/2021.11.11.468196

**Authors:** Tobias Heinen, Chen Xie, Maryam Keshavarz, Dominik Stappert, Sven Künzel, Diethard Tautz

## Abstract

*Map2k7* (synonym *Mkk7*) is a conserved regulatory kinase gene and a central component of the JNK signaling cascade with key functions during cellular differentiation. It shows complex transcription patterns and different transcript isoforms are known in the mouse (*Mus musculus*). We have previously identified a newly evolved testis specific transcript for the *Map2k7* gene in the subspecies *M. m. domesticus*. Here, we identify the new promotor that drives this transcript and find that its transcript codes for an open reading frame (ORF) of 50 amino acids. The new promotor was gained in the stem lineage of closely related mouse species, but was secondarily lost in the subspecies *M. m. musculus* and *M. m. castaneus*. A single mutation can be correlated with its transcriptional activity in *M. m. domesticus* and cell culture assays demonstrate the capability of this mutation to drive expression. A mouse knock-out line in which the promotor region of the new transcript is deleted reveals a functional contribution of the newly evolved promotor to sperm motility and to the spermatid transcriptome. Our data show that a new functional transcript (and possibly protein) can evolve within an otherwise highly conserved gene, supporting the notion of regulatory changes contributing to the emergence of evolutionary novelties.

## Introduction

Mitogen activated protein kinase (MAPK) pathways are highly conserved throughout eukaryotes and trigger multistep signaling cascades mediating transcriptional response upon reception of outside stimuli (Chang and Karin 2001, English et al. 1999). *Map2k7* belongs to the JNK group of kinases and acts as its specific activator (Fleming et al. 2000, Holland et al. 1997, Kishimoto et al. 2003, Takekawa et al. 2005, Tournier et al. 2001, Tournier et al. 1997, Wang et al. 2007). Cellular stresses like UV and gamma irradiation, osmotic shock and drug treatments on the one hand and different inflammatory cytokines, such as tumor necrosis factor, interleukin-1 or interleukin-3 on the other hand, lead to JNK pathway activation (Chang and Karin 2001, Foltz et al. 1998, Moriguchi et al. 1997, Nishina et al. 2004). Downstream targets of JNK include transcription factors (Yang et al. 2003) as well as other proteins, for example microtubule-associated proteins (Chang et al. 2003) and members of the *Bcl2* family (Deng et al. 2003, Lei et al. 2002). The JNK pathway has several known functions in the immune system, in apoptosis and in developmental processes (Dong et al. 2000, Nishina, Wada and Katada 2004, Sabapathy et al. 1999, Wada et al. 2004, Wang, Destrument and Tournier 2007). Double mutant mice lacking the JNK1 and JNK2 isoforms, as well as *Map2k7* total knockout mice, lead to embryonic lethality (Wang, Destrument and Tournier 2007)..

In their report on the first identification of *Map2k7* in mice, Tournier and colleagues (Tournier, Whitmarsh, Cavanagh, Barrett and Davis 1997) showed by Northern blotting the expression of a long transcript in various organs, plus an additional shorter transcript specific to the testis. While all the further studies concentrated on the longer transcript, the origin and function of the shorter one was left unaccounted for. In a systematic study for differentially expressed genes in mouse populations, we identified *Map2k7* to be differentially expressed in testis between wild populations of *M. m. domesticus* and *M. m. musculus*, with strongly elevated expression in *M. m. domesticus* (Harr et al. 2006). A cis-trans test via allele specific expression analysis in F1 hybrids of both subspecies demonstrated that the expression change is caused by a cis-acting sequence. It turned out that the elevated expression level can be correlated with the additional testis specific transcript in the *M. m. domesticus* subspecies which is absent in *M. m. musculus*. This suggested that a new testis-specific promotor had evolved within the *Map2k7* gene (Harr, Voolstra, Heinen, Baines, Rottscheidt, Ihle, Mueller, Bonhomme and Tautz 2006). Given the highly conserved nature of the *Map2k7* gene, such an evolution of a strong new promotor is of special interest. We present here comparative and functional data that allow inferences on the evolutionary history of the new promotor, which includes both a new origination event, as well as a secondary loss event triggered by a single mutation in one of the subspecies. The knockout analysis proofs that the new promotor has assumed a new function in the maturation of spermatids and the regulation of the transcriptome during this phase.

## Materials and Methods

### Ethics statement

The work did not involve in vivo experiments with animals. Mouse samples were taken from mice derived from the maintenance of the mouse strain collections at our institute (Harr et al. 2016). Maintenance and handling of mice in the facility were conducted in accordance with German animal welfare law (Tierschutzgesetz) and FELASA guidelines. Permits for keeping mice were obtained from the local veterinary office ‘Veterinäramt Kreis Plön’ (permit number: 1401–144/PLÖ-004697). The respective animal welfare officer at the University of Kiel was informed about the sacrifice of the animals for this study.

### In situ hybridization and Northern blotting

In situ detection of Map2k7 RNA was performed by hybridization with a digoxigenin (DIG) labeled probe (TAUTZ and PFEIFLE 1989). For probe generation, a fragment spanning Map2k7 exons 5-10 was amplified from testis C57Bl6 cDNA with primers P49 and P50 and cloned into a PCR cloning vector. The DNA fragment was reamplified from a pure plasmid clone. Reverse transcription to generate a DIG labeled probe was set up by adding 200 ng of purified PCR product to 2 μL DIG RNA Labeling Mix, 2 μL transcription buffer, 2 μL T7 polymerase, 0.5 μL RNase inhibitor (Roche, Basel). Pure water was added to the reaction mix to obtain a final volume of 20 μL. The reaction mix was incubated for 2 h at 37°C followed by a treatment with 1μL Turbo DNAse for 15 min at 37°C to remove the DNA template. The probe was precipitated with salt and alcohol, washed and re-suspended in 40 μL of 50% formamide diluted in nuclease free water (Applied Biosystems / Ambion, Austin).

All buffers and tools that were used for the following procedure were kept RNAse free. Paraffinized sections were dewaxed in xylene for 2× 10 min, washed for 5 min in ethanol, rehydrated in a series of decreasing ethanol concentration (95%, 90%, 70%, 30%; 3 min each) and washed for 5 min in PBS before postfixing them for 1 h in 4% PFA. After postfixation the tissue was washed in PBS for 2× 5 min and partially digested with 10 μg/mL proteinase K in 100 mM Tris-HCl pH 7.5 for 10 min at 37°C.

Digestion was stopped with 0.2% glycine in PBS. 2× 5 min washing in PBS was followed by 15 min incubation in 0.1 N HCl and another 2× 5 min washing in PBS was performed previous to blocking of positively charged amino acids by 0.25% acetic anhydride in 0.1 M triethanolamine pH 8.0 for 10 min. Afterwards slides were washed for 5 min in PBS and for 5 min in pure water before prehybridization for 2 h at 65°C (50% formamide, 5× SSC, 1× Denhardt’s, 0.1% Tween-20). 1 μL of DIG labeled probe was diluted in 100 μL prehybridization buffer containing 400 ng tRNA (Sigma-Aldrich, St. Louis) and denatured at 70°C for 5 min. The hybridization mix was applied to the sections and covered with coverslips. Slides were incubated over night at 65°C in a moist chamber. Next day, the sections were washed in 50% formamide containing 5× SSC and 1% SDS at 70°C for 30 min and subsequently with 50% formamide containing 2× SSC and 0.2% SDS for another 30 min at 65°C. Afterwards the sections were washed for 3× 5 min in MABS (100 mM maleic acid, 150 mM NaCl, 0.1% Tween-20 and 2 mM levamisole; adjusted to pH 7.5 with NaOH).

Samples were blocked with 1% blocking reagent (Roche, Basel) in MABS. Anti-DIG-AP antibody was applied in 1% blocking reagent in MABS by overnight incubation at 4°C. Next day, the sections were first washed 3× 10 min and then 3× 30 min in MABS. Subsequently, pH was adjusted by incubating for 3× 10 min in NTMLT buffer (100 mM Tris-HCl pH 9.5, 50 mM MgCl2, 100 mM NaCl, 100 mM levamisole, 0.1% Tween-20). BM purple solution (Roche, Basel) was applied as substrate for the alkaline phosphatase coupled with the anti-DIG antibody. Tissue was stained until the desired degree of signal was observed. Slides were washed 1 min in water and mounted with Kaisers glyceringelatine.

Detection of RNA in Northern Blotting (Alwine et al. 1977) was performed with radioactively labeled probes generated from the same clone that was used for in situ hybridization. Probes were labled with ^32^P-dCTP (Hartmann Analytic, Braunschweig) by the use of the Rediprime II DNA Labeling Kit (GE Healthcare Life Science, Little Chalfont) according to the manufacturer’s manual. Labeled probes were cleaned up with MicoSpin S-200 HR columns (GE Healthcare Life Science, Little Chalfont) according to the manufacturer’s manual.

10 μg of total RNA per sample were diluted in 15 μL nuclease free pure water (Applied Biosystems/Ambion, Austin) and mixed with 10 μL sample buffer (50% formamide, 5.18% formaldehyde, 2.5× MOPS, 0.1 mg/ml ethidiumbromide and 2.5× blue marker). Samples were heat-denatured for 5 min at 70°C and separated on an agarose gel (1.2% agarose, 0.666% formaldehyde, 1× MOPS). The RNA lanes were blotted through classic upward blot onto a Amersham Hybond N+ membrane (GE Healthcare Life Science, Little Chalfont) by neutral transfer (20× SSC) over night. Membranes were baked for 2 h at 80°C and prehybridized in ExpressHyb (Clontech, Mountain View) at 65°C for 1h. Radioactively labeled probe was added to the prehybridized blot and hybridization took place over night at 65°C in a rotating oven. Next day, the blots were washed 10 – 40 min in 2× SSC containing 0.05% SDS at RT and subsequently washed for 5 – 30 min with 0.1× SSC containing 0.1% SDS at 50°C. After washing, the blots were dipped in 2× SSC, sealed in a plastic bag and analyzed via autoradiography using Kodak Biomax-MS films (Kodak, Rochester).

### Promotor tests in cell culture

The promotor tests in cell culture required to set up an appropriate expression system. The details on the construction and testing of this system are described in (Heinen 2008). It resulted in the construction of a “Luciflip plasmid” that contains the following elements in the given order: PGK-promoter, ATG, FRT, splice acceptor, double polyA signal, EcoRI site, Kozak, Luc-MYC, intron, polyA signal and was the basis for the Map2k7 alpha reporter assay. For this, fragments spanning -487 to +43 relative to the transcription start of the Map2k7 alpha promoter were amplified from genomic DNA of *M. m. musculus* and *M. m. domesticus* using the primer pair P318 / P319. A two-step PCR strategy was pursued to generate fragments with a deleted insulator motive. Two separate PCRs with the primer pairs P318 / P320 and P321 / P319 were run on top of the cloned promoter fragment. The primers P319 and P320 bind just right upstream and downstream of the insulator sequence and were tailed with a sequence stretch which is homologous to the sequence on the opposite part exactly beyond the insulator. The other primer is one of the primers that were used in the first PCR. Thus, the promoter fragment is divided into two fragments each defined by an inner and an outer primer. The inner edges overlapped, but were lacking the insulator. Both PCR products were cleaned up and included into another PCR without primers. After 5 cycles, the outer primers P318 and P319 were added to the reaction and PCR continued as usual. The resulting product was cloned into a PCR cloning vector and sequenced with M13 primers.

The second version of all 4 fragments has an additional upstream CMV enhancer. For this, the CMV enhancer was amplified with the primers P322 and P323 using the phrGFPII-1 plasmid (Agilent Technologies, Santa Clara) as template. The resulting product was cloned into a PCR cloning vector and validated by M13 primer sequencing. Both CMV primers and the upstream primer that was used for promoter fragment generation (P318) were tailed with an XhoI restriction site overhang. CMV enhancer fragments were retrieved by XhoI digestion and ligated into the XhoI site of all 4 Map2k7 promoter fragments. The assembled fragments were cut out by EcoRI digestion and cloned into the EcoRI site of Luciflip. The orientation of the inserts was controlled with an XhoI digest. All 8 Luci-flip constructs were sequenced with the primers P293, P358 and P199 to control the inserts. No mutations were found.

The 8 expression constructs were transfected into NIH/3T3 cells. For this, NIH/3T3 fibroblast cells were grown in DMEM medium containing sodium pyruvate, non-essential amino acids, L-glutamine penicillin/streptomycin (all from Invitrogen, Carlsbad) and 10% fetal calf serum (PAN, Aidenbach) at 37°C incubation maintaining 5% CO_2_ concentration. One day before transfection, 3 × 10^3^ NIH/3T3 cells were seeded with 70 μL medium into each well of 96-well plates and grown over night. Cells were co-transfected with Luciflip plasmid and a pGL3 plasmid (Promega, Mannheim) which contains firefly luciferase under the control of SV40 promoter. Firefly luciferase was used to normalize transfection efficiency. Therefore, 0.18 μL Fugene 6 reagent (Roche, Basel) was added to 4.82 μL serum free medium and incubated for 5 min at RT. Subsequently, 30 ng of Luciflip and 30 ng of pGL3 DNA were mixed and added to the medium containing Fugene 6. The mixture was incubated for 20 min and added to one well of the 96 well plate containing NIH/3T3. The transfection was performed for the Luciflip-CMV construct and for an empty Luciflip plasmid as blank control. Each transfection was performed in 8 replicates in parallel. Cells were incubated over night. Next day, firefly and renilla luciferase substrates were applied using the Dual-Glo Luciferase Assay System (Promega, Mannheim) according to the manufacturer’s manual. Relative light units were measured with a Mithras LB 940 Luminometer (Berthold Technologies, Bad Wildbad). For every well, the renilla luciferase signal was divided by the firefly luciferase signal to normalize transfection efficiency. For every construct, median and standard deviation was calculated from the 8 individual replicates.

### Construction of knockout mice

The general scheme for the construction of the promotor knockout is shown in suppl. file S2. It consisted two steps. In the first step, in neomycin cassette was inserted at position chr8:4,239,573 (mm10) via homologous recombination in embryonic stem cells, whereby 593 bp of the promotor region, including the start sites, was deleted. These cells were then used to generate transgenic mice via injection into blastocycsts. In the second step, the neomycin cassette was removed via flp recombination at the FRT sites in the mice. The annotated wildtype sequence in this region, as well as the neomycin cassette and the final sequence after recombination are given in suppl file S2. The generation of the KO mice was done by inGenious Targeting Laboratory (iTL), Stony Brook. The mice were then transferred into our facility and backcrossed against C57Bl6/J until final analysis after about 15 generations.

### Sperm analysis

Testis and epididymis were dissected from the right side of each mouse and weight was measured. The cauda epididymis was excised, immediately transferred in 250 μl human tubular fluid medium (Millipore, Billerica), punctured with a needle and placed at 37°C, 5% CO_2_ for 20 minutes. After 5 min of incubating a 6 μl drop of medium with dispersed spermatozoa was transferred onto a warmed glass slide and covered with a 20 × 20 mm coverslip. Progressive motility was estimated by phase contrast microscopy at a magnification of x 200 according to WHO (World Health Organization, 1992) in a Neubauer chamber. At least 200 sperm cells (but up to 700) in different areas of the slide were counted per animal and the percentage of different classes of motility was calculated per animal.

### RNA-Seq and data analysis

The testis tissues of 8 WT and 8 knockout mice were carefully collected and immediately frozen in liquid nitrogen. Total RNA was purified using QIAGEN RNeasy Microarray Tissue Mini Kit (Catalog no. 73304), and prepared using Illumina TruSeq Stranded mRNA HT Library Prep Kit (Catalog no. RS-122–2103), and sequenced using Illumina NextSeq 500 and NextSeq 500/ 550 High Output v2 Kit (150 cycles) (Catalog no. FC-404–2002). Raw reads in FASTQ format were trimmed with Trimmomatic (0.38) (Bolger et al. 2014), and only the reads left in pairs were used for further analysis. The trimmed reads were mapped to the mouse reference genome GRCm39 (Howe et al. 2021, Waterston et al. 2002) with HISAT2 (2.2.1) (Kim et al. 2015) and SAMtools (1.9) (Li et al. 2009), and the mouse gene annotation in Ensembl (Version 104) was used for indexing the genome, i.e., the options “--ss” and “--exon” were used for command “hisat2-build”. The numbers of fragments uniquely mapped to the genes annotated in Ensembl (Version 104) were calculated with featureCounts (2.0.3) (Liao et al. 2014). Principle component analysis on variance stabilizing transformed fragment counts and differential expression analysis on raw counts were performed with DESeq2 (1.30.1) (Love et al. 2014, Zhu et al. 2019).

### Primer list

All primers were obtained from Metabion (Martinsried). Sequences are listed 5’> 3’.

P49 AATTAACCCTCACTAAAGGGGAGCATCGAGATTGACCAGA

P50 TAATACGACTCACTATAGGGGCTCGGATGTCATAGTCAGG

P318 CTCGAGTGACCAACTACTTTTCACTATTGCTG

P319 CAAGCTGTGAAGGTCAGTCAGG

P320 TGGTGGACAAGCTGGATCTAGAAAGGAAGAGGAAGCACT

P321 CTCTTCCTTTCTAGATCCAGCTTGTCCACCATGACC

P322 CTCGAGCGCGTTACATAACTTACGGTAAA

P323 CTCGAGCAAAACAAACTCCCATTGACG

## Results

Comparison of testis expression of *Map2k7* in two mouse subspecies (*Mus musculus domesticus*, and *Mus musculus musculus*) via Northern blotting and in situ hybridization shows a major difference between the two subspecies (Figure 1). We included in our analysis testis samples from the inbred strain C57Bl6/J (derived from *M. m. domesticus* (Frazer et al. 2*007)*), as well as from wild caught mice that were kept under outbreeding conditions (Harr, Karakoc, Neme, Teschke, Pfeifle, Pezer, Babiker, Linnenbrink, Montero, Scavetta, Abai, Molins, Schlegel, Ulrich, Altmuller, Franitza, Buntge, Kunzel and Tautz 2016). Northern blotting revealed a weak 3.5 kb and a strong 1.6 kb band in *M. m. domesticus* (Figure 1A), but only the 3.5kb band in *M. m. musculus*, with the 1.6kb band completely missing. For the inbred strain this confirms the previous observations by (Tournier, Whitmarsh, Cavanagh, Barrett and Davis 1997) and for the wild type strains the observation by (Harr, Voolstra, Heinen, Baines, Rottscheidt, Ihle, Mueller, Bonhomme and Tautz 2006).

**Figure 1:**
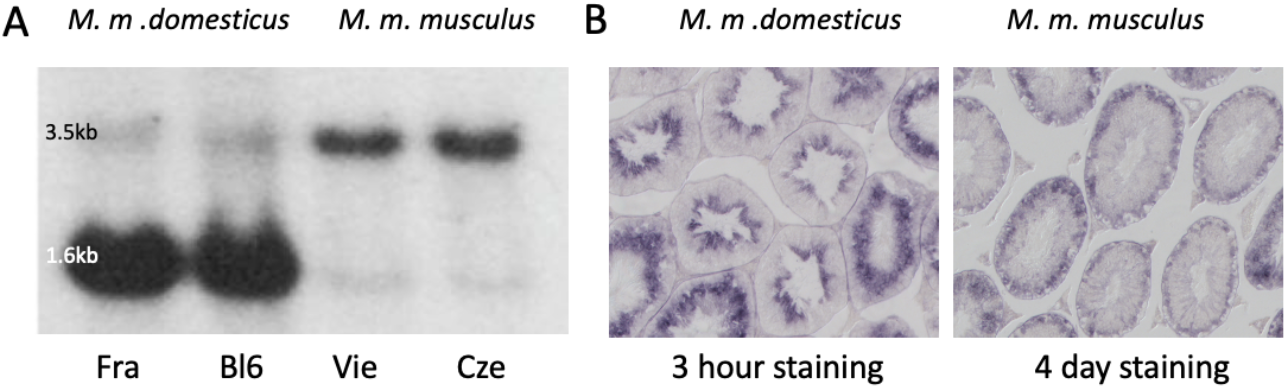
*Map2k7* expression in *M. m. domesticus* versus *M. m. musculus*. (A) Northern blots with testis RNA from individuals of a wild population from France (Fra), the laboratory inbred strain C57Bl6/J (Bl6), a wild caught population from Vienna (Vie) and a wild caught population from the Czech Republic (Cze). The strongly expressed 1.6kb band is only visible in the two individuals from the *M. m. domesticus* sub-species. (B) In situ hybridization with the *Map2k7* probe on cross sections of seminiferous tubules of an individual from *M. m. domesticus* (left) and an individual from *M. m. musculus* (right). Note that the left signal developed already after 3 hours of color incubation, while the right signal developed only after 4 days of color incubation.

To assess the stage of spermatid development at which *Map2k7* is expressed, we used in situ hybridization on testis sections from *M. m. domesticus* and *M. m. musculus*. Testis tissue mainly consists of seminiferous tubules, which are the location of spermatogenesis. Spermatogonial stem cells adjacent to the inner tubule wall divide and form spermatocytes which undergo meiosis. After meiosis the spermatocytes develop into spermatids and change morphologically from round spermatids to elongated spermatids before the generation of mature spermatozoa is completed. The three main stages, spermatogonia, spermatocytes and spermatids, are classified into further substages (Russell et al. 1990). Sperm precursor cells are embedded in Sertoli cells which define the shape of the spermatogenic epithelium and support the germ cells. Through the influence of Sertoli cells, developing sperm precursor cells proceed towards the lumen of seminiferous tubules according to their degree of maturation. Terminal spermiation releases the sperm cells into the luminar fluid of the tubules that transfers them to the epididymis. Hence, ring-shaped zones representing different cell stages can be distinguished in a transverse section of seminiferous tubules.

In situ hybridization results on testis sections using the same probe that had been used for the Northern blotting are shown in Figure 1B. In *M. m. domesticus*, we find a strong signal in post-meiotic spermatid stages. In contrast, *Map2k7* expression pattern in *M. m. musculus* is very weak and becomes only visible after several days of color development. This signal is restricted to earlier stages and we interpret it as the expression of the long transcript. This expression would be expected to be present also in *M. m. domesticus*, but the sections are over-stained after several days of incubation, making it impossible to visualize this directly.

To better resolve the transcript structures and to assess the origin of the new promotor, we made use of the transcriptome data described in (Harr, Karakoc, Neme, Teschke, Pfeifle, Pezer, Babiker, Linnenbrink, Montero, Scavetta, Abai, Molins, Schlegel, Ulrich, Altmuller, Franitza, Buntge, Kunzel and Tautz 2016). Figure 2 shows the read coverage as browser tracks aligned to the annotated versions of the *Map2k7* transcripts in the UCSC browser. Note that there are two versions of the 3.5kb transcript, differing by the inclusion/exclusion of a small exon. Further, there are four versions of the 1.6kb transcript, whereby only two are due to the new promotor, while the other ones are splice variants of the longer transcript (Figure 2 - top panel). The set of browser tracks representing the transcriptomes from 10 different tissues of individuals from a *M. m. domesticus* population show that the new exon (highlighted in grey) is only expressed in the testis (Figure 2 - middle panel). The second set of browser tracks shows testis transcriptomes from different populations of the sub-species, as well as closely related species. The new exon is only present in *M. m. domesticus* populations (DOM), as well as in the sister species *M. spretus* and *M. spicilegus*, but absent in the subspecies *M. m. musculus, M. m. castaneus* as well as the further distant mouse species *M. matheyi* and the wood mouse *Apodemus* (Figure 2 - lower panel).

**Figure 2:**
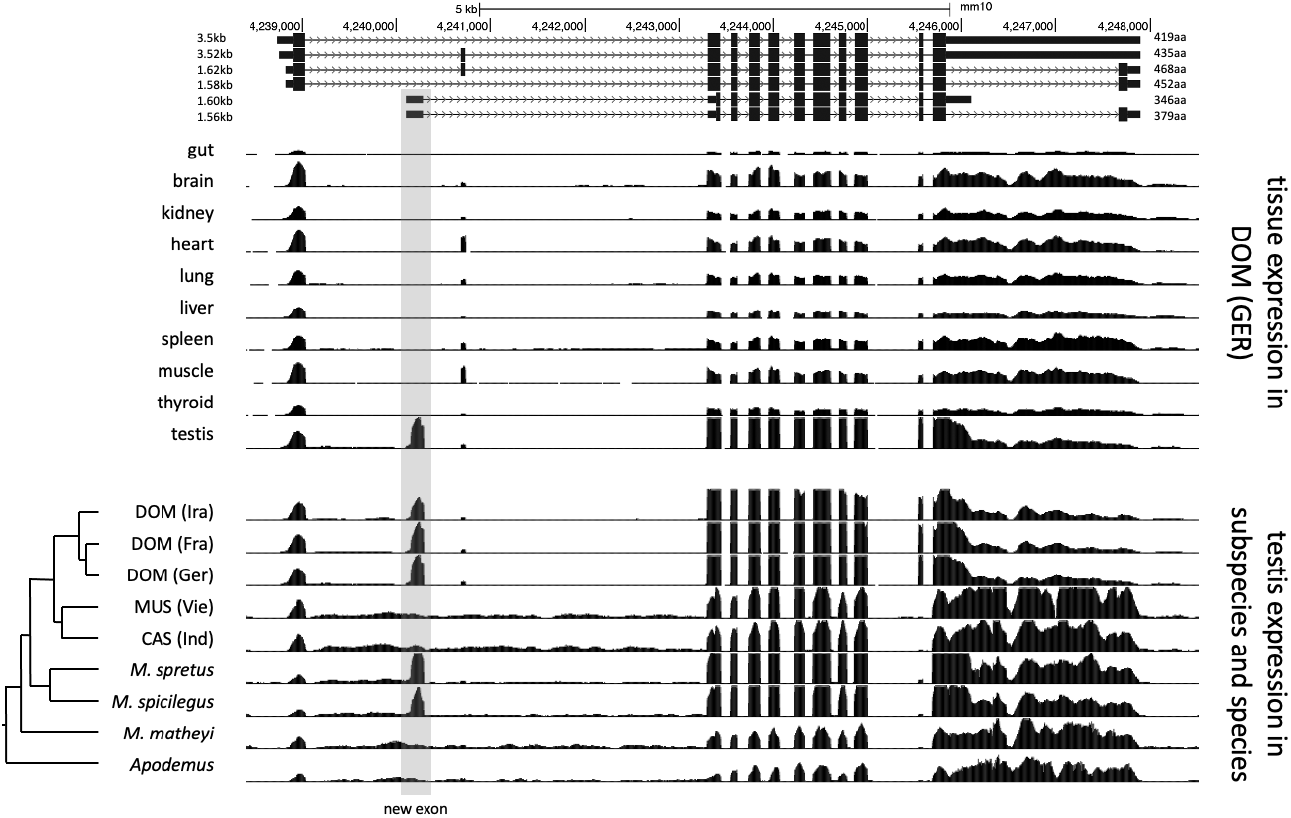
*Map2k7* transcript variants and transcriptome read coverage. The figure is based on UCSC genome browser tracks. The top panel shows the different transcript and splice versions from the mouse reference genome, which reflects *M. m. domesticus* and therefore includes the new promotor/exon (highlighted by grey shading). The middle panel is based on data from (Harr, Karakoc, Neme, Teschke, Pfeifle, Pezer, Babiker, Linnenbrink, Montero, Scavetta, Abai, Molins, Schlegel, Ulrich, Altmuller, Franitza, Buntge, Kunzel and Tautz 2016) and shows transcriptome read mapping tracks for different tissues. The lower panel is based on transcriptome data from (Harr, Karakoc, Neme, Teschke, Pfeifle, Pezer, Babiker, Linnenbrink, Montero, Scavetta, Abai, Molins, Schlegel, Ulrich, Altmuller, Franitza, Buntge, Kunzel and Tautz 2016) and (Neme and Tautz 2016) and shows transcriptome read mapping tracks for different populations, sub-species and species. The phylogenetic relationships are depicted to the left. DOM: *M. m. domestics*, MUS: *M. m. musculus*, CAS: *M. m. castaneus*.

Tournier and colleagues have earlier identified the different isoforms of *Map2k7* by screening testis cDNA clones from laboratory mice and implemented a nomenclature (Tournier et al. 1999). Consistent with the data described above, they found three different 5’-versions (named α, β and γ isoforms) and two 3’-variants (named 1 and 2 isoforms) whose combinations are obtained by alternative splicing. The nomenclature can be complemented by a 3’-variant named 3 that defines the long ∼3.5 kb transcript. Northern blotting and qPCR using different exon-specific probes and primers (Suppl. file S1) lead us to the conclusion that the highly expressed testis specific ∼1.6 kb fragment corresponds to the transcript that was called Mkk7-α1 in (Tournier, Whitmarsh, Cavanagh, Barrett and Davis 1999) and that we will call Map2k7-α1 in the following to account for the change in the official gene nomenclature.

The transcripts Map2k7-β3 and Map2k7-γ3 (γ includes an additional small exon without an alteration of the reading frame), include the JNK-binding site (D-domains) in the first exon (Figure 3). They are about ∼3.5 kb in size and can be found in all analyzed populations and species. The newly evolved transcription start is situated within the first intron of the conserved transcripts. Its transcript does not include the exon with the D-domains, but encodes potentially a protein with the kinase and DVD domain. (Tournier, Whitmarsh, Cavanagh, Barrett and Davis 1999) have shown that this truncated protein has a detectable, but very weak kinase activity when expressed from an expression vector in cell culture. However, the first AUG in the new transcript is before this long ORF and in a different reading frame. It codes for a novel 50aa protein (Figure 3) and the nucleotides surrounding the start codon of this new ORF (UGGCCAACG <u>AUG</u> G) match much better to the Kozak-consensus-sequence (Kozak 1987) than the nucleotides surrounding the start codon of the remaining reading frame of the Map2k7-α1 transcript (CCCCGCCAC <u>AUG</u> C). A purine at position -3 and a guanine at position +4 are the most important sequence elements for the initiation of translation. It is therefore questionable whether the shortened form of *Map2k7* is translated at all under natural conditions.

**Figure 3:**
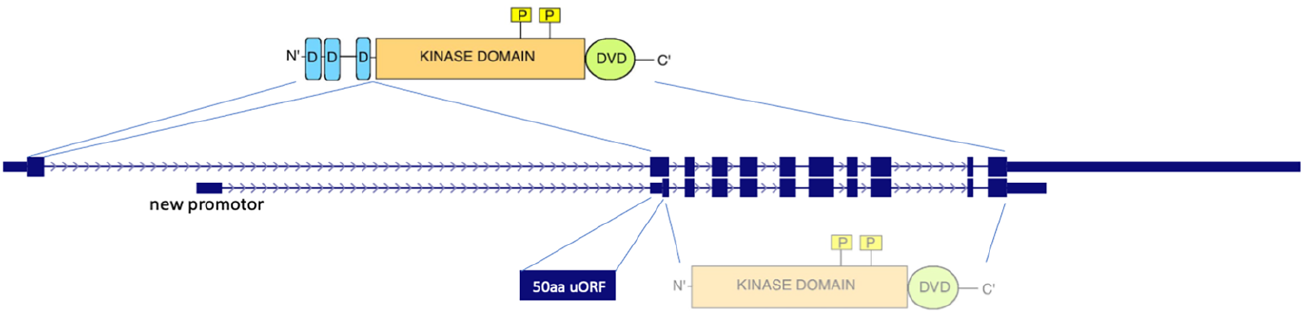
Comparison of the conserved and the new transcripts and their coding potentials. Transcript depictions are taken from the UCSC browser annotations (see also Figure 2), whereby only the two main transcripts are shown, the conserved one (upper) and the new one starting from the new promotor (lower). The protein domains of the *Map2k7* functional kinase are depicted on the top, including the JNK-binding sites (D-domains in blue), the kinase domain (in orange) with its functional phosphorylation sites (yellow), as well as the C-terminal DVD domain (green). The first AUG in the new transcript is in the second exon and it would result in a 50aa protein, which represents a *de novo* generated protein without known domain. If this AUG would not be used, the next AUG in a different frame could potentially lead to the expression of a truncated version of the *Map2k7* protein, containing only the kinase and the DVD domains.

Based on the phylogenetic tree of wild mice (Chevret et al. 2005, Guenet and Bonhomme 2003), we can infer that the new 1.6 kb transcript has arisen before the branching of *M. spicilegus* and *M. spretus* at least 2 million years ago, but not more than 6 million years ago, since it is not present in the outgroups *M. matheyi* and *Apodemus* (Figure 2). Note that while these outgroups show some general level of intron transcription in the respective area, this does not reflect the clear exon structure that is seen in the other species. The new promotor has secondarily disappeared in the lineage of *M. m. musculus* and *M. m. castaneus*, apparently because of a crucial T/A substitution (see below).

### Promotor analysis

To further understand the cis regulation of the spermatid specific 1.6 kb Map2k7α1 transcript, we analyzed genomic sequences in a 500 bp window upstream of the transcription start site in different wild populations, subspecies and species of *M. m. domesticus, M. m. musculus, M. m. castaneus, M. spretus and M. spicilegus* (Figure 4). Only one SNP at -84 bp (with respect to the start site of transcription - see also legend Figure 4 for alternative start sites) is correlated with the expression of the spermatid specific isoform in the different populations. An adenine is found in populations with expression of the new transcript, whereas thymine is present in those without expression (Figure 4).

**Figure 4:**
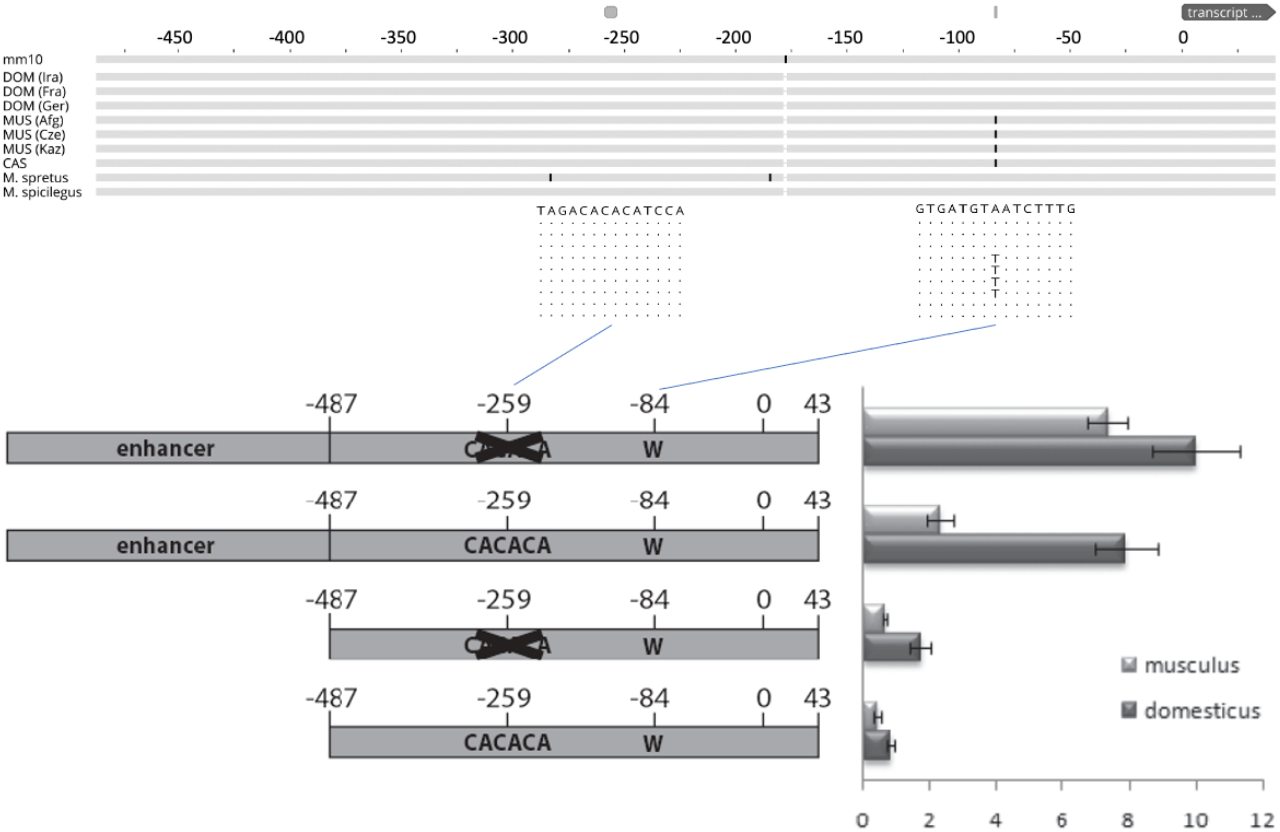
Functional test of the promotor region driving the new transcript. The top shows the alignment of genomic sequences from populations, sub-species and species based on the genome data from (Harr, Karakoc, Neme, Teschke, Pfeifle, Pezer, Babiker, Linnenbrink, Montero, Scavetta, Abai, Molins, Schlegel, Ulrich, Altmuller, Franitza, Buntge, Kunzel and Tautz 2016), aligned to the mouse mm10 reference sequence. The fragment shown represents the one used for the promotor studies - only replacements with respect to the reference are marked, the two relevant regions discussed in the text are enlarged with their respective sequences. DOM represents *M. m. domesticus* populations, MUS represents *M. m. musculus* populations, CAS represents a *M. m. castaneus* population. T/A is the only mutation that correlates with the expression of the new promotor. Note that the transcriptional start site, marked as “0”, is located between the annotated site for this transcript, which would be 31bp further upstream and the site from which the bulk of the transcripts generated in the RNASeq experiment starts, which is 8bp downstream (see suppl. file S2 for a corresponding sequence depiction of this region). The bottom shows the scheme of the four constructs that were tested in cell culture, as well as their expression levels measured as fluorescence intensity (see Methods). Error bars indicate standard deviations from eight replicates each.

Empirical data provide evidence that specific transcription activity in reproductive tissues and particularly in spermatogenesis is regulated by very short proximal promoters (Blaise et al. 2001, Han et al. 2004, Li et al. 1998, Reddi et al. 1999, Scieglinska et al. 2004, Topaloglu et al. 2001, Zambrowicz et al. 1993). Those studies demonstrated that proximal promoters shorter than 300bp, or even less than 100bp, are sufficient to drive spermatid specific expression in mice. Unique mechanisms of gene regulation are postulated to exist in post-meiotic cells (Acharya et al. 2006, Somboonthum et al. 2005). Additionally, it was shown that a 5’- CACACA motive ∼170 bp upstream of the transcription start serves as an insulator in the spermatid specific expression of the SP-10 gene (acrosomal vesicle protein 1 - Acrv1) and it was suggested that insulators might generally play an important role in maintaining spermatid specific transcription (Abhyankar et al. 2007, Acharya, Govind, Shore, Stoler and Reddi 2006, Reddi et al. 2003, Reddi et al. 2007).

Such a 5’- CACACA motif can be found at around -259 base pairs upstream the transcription start site of the testis specific Map2k7-α1 RNA (Figure 4). These considerations raise the question, whether a short sequence carrying the -84 A mutation in combination with the -259 5’- CACACA motif would meet the requirements to serve as a testis specific promoter. Therefore, we tested a fragment representing the genomic sequence between -487/+43 of the new *Map2k7* testis promoter in cell culture-based luciferase expression assays (Figure 4).

The expression of most interest in this context is restricted to late spermatids. Culturing this type of cells is very difficult due to its haploid post meiotic stage with condensed chromatin. A well-established spermatid cell culture model is not available and alternative cell lines have the disadvantage that they will most likely not recognize the spermatid specific *Map2k7* promotor. In the absence of better options, we have chosen the widely used NIH/3T3 fibroblast cell line for this experiment.

It cannot be expected that the tested fragment is sufficient to drive luciferase expression in non-spermatid cells, but it is likely that the expression level can be raised by deleting the 5’- CACACA motive at -259, if the assumption is correct, that this sequence maintains spermatid specific transcription by acting as an insulator in other cells. Thus, -487/+43 fragments lacking the 5’-CACACA motive at -259 were generated as well. It can be assumed that the -487/+43 fragments do not contain enhancer elements which promote expression in fibroblasts. Therefore, a CMV enhancer was ligated upstream to both versions. All four constructs (wild type, wild type with deleted insulator, CMV enhancer + wild type, CMV enhancer + wild type with deleted insulator) were created as *M. m. domesticus* variants with an adenine at position - 84 and as *M. m. musculus* variants with a thymine at position -84. We find that the *M. m. domesticus* variant generates significantly higher signals compared to *M. m. musculus* in every combination. The different replicates are consistent, indicated by relatively small standard deviations (Figure 4). Deletion of the 5’-CACACA motive at -259 indeed increases the expression strength in both variants. The presence of an enhancer potentiates the effects as expected. These data provide strong evidence that the adenine at position -84 enhances the activity of the basal promotor. For this reason, it can be supposed that a major contribution of this mutation to the expression difference between *M. m. domesticus* and *M. m. musculus* in late spermatids is likely. The sequence 5’-CACACA represses the action of an adjacent enhancer to a certain extent. This finding supports the hypothesis that it acts as an insulator in the spermatid specific *Map2k7* promotor.

### Functional analysis

To assess a possible functional role of the new spermatid specific *Map2k7* promotor, a knock-out mouse was generated in which the promotor was deleted in the *M. m. domesticus* background (see Methods and targeting strategy in suppl. file S2). The knock-out was designed in a way that it should not interfere with the conserved *Map2k7* transcript.

The knockout animals were fully viable and showed no overt phenotype. KO animals are on average a bit heavier, but have lower normalized testis weights (Table 1). Given the specific expression of the new transcripts during a crucial phase of sperm maturation, we assessed also sperm motility phenotypes. KO animals have fewer motile sperm and fewer progressive sperm (Table 1). All differences are significant at p< 0.05 (t-test, 2 sided).

**Table 1.** Sperm phenotypes.

| genotype |  | mouse weight<br>(g) | normalized<br>testis weight<br>(g) | motile sperm<br>(%) | progressive<br>sperm<br>(%) |
| --- | --- | --- | --- | --- | --- |
| WT (N=10) | averages: | 25.14 | 0.20 | 48 | 26 |
|  | *SD*: | *1.42* | *0.03* | *6* | *5* |
| KO (N=18) | averages: | 27.59 | 0.17 | 42 | 22 |
|  | *SD*: | *2.07* | *0.01* | *7* | *5* |
|  | P-values: | 0.003 | 0.001 | 0.021 | 0.031 |

### RNASeq analysis

A comparative RNASeq analysis with RNA from knock-out mice versus wild type mice was used to find out whether the 1.6 kb Map2k7-α1 testis specific transcript is influencing the expression of other genes. The RNA was collected from three different tissues of the male reproductive organs, the testis, the caput epididymis and the cauda with eight biological replicates each. The testis is the place of the primary sperm production. The sperm from the testis move through the caput epididymis where they mature and are eventually stored in the cauda. While the chromatin of post meiotic sperm is condensed, there is still some transcriptome turnover (Ren et al. 2017) and the epididymal cells contribute to this transcriptome turnover as well (Shi et al. 2021).

The overall analysis of the transcriptome data in the PCA analysis shows that the samples from each of the three tissues are very different, implying that there is indeed a major turn-over of RNA between these stages, either due to differential stability, or new transcription. On the other hand, differences between wild type and knockout are much smaller (Figure 5A). Still, since we used eight replicates for each tissue, we have a very high sensitivity to detect even small transcriptome changes (Xie et al. 2020). Accordingly, we find thousands of genes with significant expression differences (i.e., p_adj_ values <0.05 in the DSeq2 analysis), but mostly with relatively low log2fold-changes (Figure 5B-D). Interestingly, however, the cauda samples show a set of genes with very high positive log2fold changes (see set of dots on the top of the panel for “cauda in Figure 5D).

**Figure 5:**
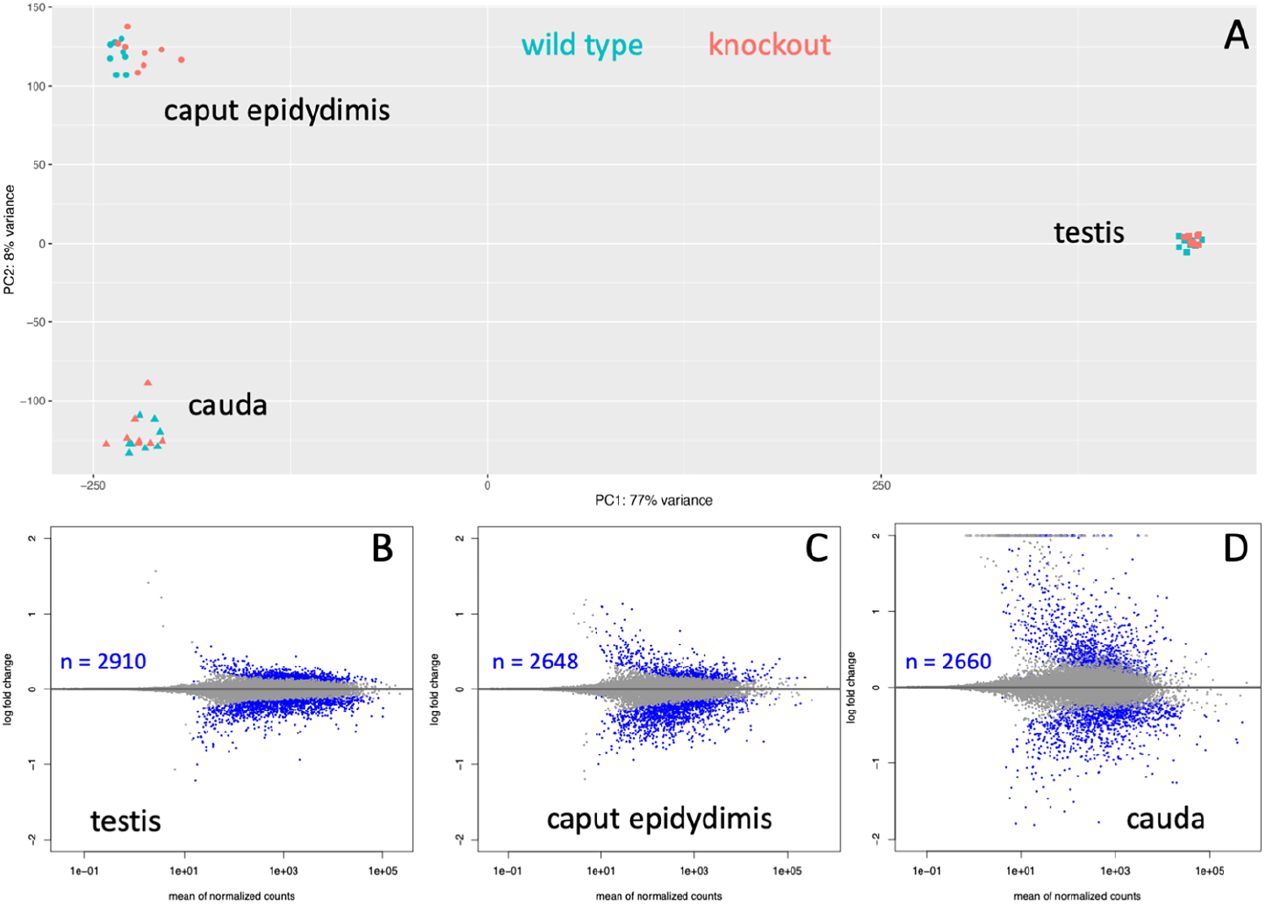
Whole transcriptomes analysis of three tissues from wild type versus knockout mice. (A) Overall PCA comparison. Strong differentiation is seen between the tissue samples, implying that that transcript sets are very different. The differences between wild type and knouts are much smaller for each tissue. (B-D) Significantly differentially expressed genes. Each gene is represented by a dot, genes with padj < 0.05 values are plotted as blue dots. The number of significantly differentially expressed genes is provided as inset for each tissue.

First, we asked whether kinase signaling processes are specifically affected, which would suggest that the shortened protein of the Map2k7-α1 transcript could be involved in signaling. However, the top biological process GO terms among the significant genes do not include “kinase signaling”, “signal transduction” or “Jnk cascade” in either of the data sets (based on a GO analysis with Panther (Mi et al. 2013) - see suppl Table 1). Instead, the top GO terms indicate an involvement in meiotic division and chromosome segregation for testis, an involvement in extracellular matrix organization for the caput epididymis and an involvement in peptide biosynthetic processes for the cauda (suppl Table 1b, d, f). Hence, it is unlikely that the primary function of the Map2k7-α1 transcript is related to residual kinase signaling activity.

To get a further insight into the functional changes in the knockout animals, we focused on the genes that are most highly expressed in the respective tissues (based on the length-normalized baseMean counts of the RNASeq data), since small concentration changes in such genes could have a more marked influence on the phenotype.

For most of the highly expressed genes that we identify in the significant gene lists for testis and cauda epididymis, one can retrieve functional information from knockout experiments in mice and almost all of these find an effect on sperm maturation and/or sperm mobility (Table 2). Hence, while the relative expression changes are not large, it is well possible that the effect of the Map2k7-α1 transcript knockout is mediated via these genes. Interestingly, for the cauda, we find a rather different pattern. Most of the top expressed genes in the list are not specific for the cauda, but are more broadly expressed (e.g., an enzyme, actin and a ribosomal protein). Interestingly, several code for immunity proteins, including sperm-specific antimicrobial peptides. Another major difference in the cauda transcriptome is a set of 37 genes with very high log2fold changes, i.e., expression at a much higher level in the knockout than in the wild type. Intriguingly, most of these are generally expressed motor proteins and the role of such proteins in spermiogenesis has only recently been fully recognized (Wu et al. 2021).

**Table 2.** List of top significant genes in the RNASeq analysis.

| tissue | gene name | description | log2 fold-change | padj | expression | function | literature |
| --- | --- | --- | --- | --- | --- | --- | --- |
| *testis (top 10 most highly expressed genes that are specifically expressed in testis)<sup>#1</sup>* |  |  |  |  |  |  |  |
|  | *Tnp2* | Nuclear transition protein 2 | -0.37 | 0.0021 | testis | conversion of nucleosomal chromatin in spermatids, involved in sperm motility | (Adham et al. 2001) |
|  | *Smcp* | Sperm mitochondrial-associated cysteine-rich protein | -0.25 | 0.0006 | testis | involved in sperm motility and fertilization | (Nayernia et al. 2002) |
|  | *Gapdhs* | glyceraldehyde-3-phosphate dehydrogenase, spermatogenic | -0.12 | 0.0206 | testis | required for normal sperm motility and male fertility, | (Huang et al. 2017) |
|  | *Fhl4* | Four and a half LIM domains 4 | 0.09 | 0.0186 | testis | may affect sperm maturation and morphology | (Lardenois et al. 2009) |
|  | *Tuba3a* | tubulin, alpha 3A, spermatogenic | -0.14 | 0.0027 | testis | required for the production of normal spermatozoa. | (Akter et al. 2021) |
|  | *Crisp2* | Cysteine-rich secretory protein 2 | 0.10 | 0.0444 | testis | regulates calcium fluxes during sperm capacitation, essential for fertility | (Curci et al. 2020) |
|  | *Akap4* | A-kinase anchor protein 4 | 0.13 | 0.0037 | testis | major structural component of sperm fibrous sheath, plays a role in sperm motility | (Zhang et al. 2021) |
|  | *Tuba3b* | tubulin, alpha 3B, spermatogenic | -0.11 | 0.0036 | testis | required for the production of normal spermatozoa. | (Akter, Hada, Shikata, Watanabe, Ogura and Matoba 2021) |
|  | *Cox8c* | cytochrome c oxidase subunit 8C | -0.26 | 0.0000 | testis | required for the production of normal spermatozoa. | (Akter, Hada, Shikata, Watanabe, Ogura and Matoba 2021) |
|  | *H1fnt* | testis specific H1.7 linker histone | -0.15 | 0.0013 | testis | required for cell restructuring and DNA condensation during spermiogenesis | (Martianov et al. 2005) |
| *caput epididymis (top 10 most highly expressed genes that are specifically expressed in testis/epididymis)<sup>#2</sup>* |  |  |  |  |  |  |  |
|  | *Lcn8* | Epididymal-specific lipocalin-8 | 0.15 | 0.0493 | epididymis | involved in sperm maturation and motility | (Wen et al. 2021) |
|  | *Lcn9* | lipocalin 9 | 0.14 | 0.0450 | epididymis | may act redundantly to Lcn8 | (Wen, Liu, Zhu, Sun, Xiao, Lin, Zhang, Ye and Gao 2021) |
|  | *Epp13* | Epididymal protein 13 | -0.13 | 0.0378 | epididymis | no fecundity defect in knockout mice | (Noda et al. 2019) |
|  | *Cst8* | cystatin 8 (cystatin-related epididymal spermatogenic) | 0.24 | 0.0016 | epididymis | capacitation of spermatozoa | (Chau and Cornwall 2011) |
|  | *Rnase10* | Ribonuclease-like protein 10 | 0.40 | 0.0000 | epididymis | required for post-testicular sperm maturation | (Krutskikh et al. 2012) |
|  | *Adam7* | Disintegrin and metalloproteinase domain-containing protein 7 | 0.15 | 0.0108 | epididymis | required for epididymal integrity, sperm morphology and motility | (Choi et al. 2015) |
|  | *Teddm1a* | Transmembrane epididymal family member 1a | 0.28 | 0.0035 | epididymis | no information on sperm effects |  |
|  | *Teddm2* | Transmembrane epididymal family member 2 | 0.34 | 0.0001 | epididymis | no information on sperm effects |  |
|  | *Spink12* | serine peptidase inhibitor, Kazal type 12 | -0.31 | 0.0003 | epididymis | part of Spink family with redundant functions in sperm maturation | (Jeong et al. 2019) |
|  | *Defb12* | defensin beta 12 | 0.20 | 0.0033 | epididymis | immunity protein |  |

| *cauda (top 10 most highly expressed genes)<sup>#3</sup>* |  |  |  |  |  |  |  |
| --- | --- | --- | --- | --- | --- | --- | --- |
|  | *Crisp1* | Cysteine-rich secretory protein 1 | -0.46 | 0.0013 | testis, epididymis | required to optimize sperm flagellum waveform | (Gaikwad et al. 2021) |
|  | *Cd52* | CAMPATH-1 antigen | -0.69 | 0.0005 | broad | immunity protein | (Yamaguchi et al. 2008) |
|  | *Gpx3* | Glutathione peroxidase 3 | -0.73 | 0.0043 | broad | protects cells from oxidative damage, involved in the maturation of sperm cells | (Noblanc et al. 2011) |
|  | *Defb28* | Defensin beta 28 | -0.37 | 0.0404 | testis, epididymis | immunity protein |  |
|  | *Spink8* | Serine protease inhibitor Kazal-type 8 | -0.44 | 0.0009 | testis, epididymis | part of Spink family with redundant functions in sperm maturation | (Jeong, Lee, Kim, Hong, Kim, Choi, Cho and Cho 2019) |
|  | *B2m* | Beta-2-microglobulin | -0.39 | 0.0035 | broad | component of the class I major histocompatibility complex (MHC) |  |
|  | *Wfdc15b* | WAP four-disulfide core domain protein 15B | -0.75 | 0.0002 | testis, epididymis | Wfdc family members act in innate immune responses during epididymitis | (Andrade et al. 2021) |
|  | *Defb2* | defensin beta 2 | -0.38 | 0.0336 | testis, epididymis | immunity protein |  |
|  | *Acta2* | Actin, aortic smooth muscle | 0.44 | 0.0409 | broad | cell motility, may affect contractability of seminiferous tubules | (Uchida et al. 2020) |
|  | *Rplp1* | ribosomal protein, large, P1 | -0.54 | 0.0110 | broad | ribosomal protein |  |
| *cauda (top 10 genes with highest log2fold changes)<sup>#3</sup>* |  |  |  |  |  |  |  |
|  | *Pvalb* | parvalbumin | 8.93 | 0.0158 | broad | no information on sperm effects |  |
|  | *Myl2* | myosin, light polypeptide 2, regulatory, cardiac, slow | 8.36 | 0.0338 | broad | motorprotein involved in spermiogenesis | (Wu et al. 2021) |
|  | *Myh7* | myosin, heavy polypeptide 7, cardiac muscle, beta | 8.04 | 0.0405 | broad | motorprotein involved in spermiogenesis | (Wu et al. 2021) |
|  | *Myl1* | myosin, light polypeptide 1 | 7.26 | 0.0170 | broad | motorprotein involved in spermiogenesis | (Wu et al. 2021) |
|  | *Atp2a1* | ATPase, Ca<sup>++</sup> transporting, cardiac muscle, fast twitch 1 | 7.05 | 0.0008 | broad | Ca-gradient regulation, male fertility | (Prasad et al. 2004) |
|  | *Myh1* | myosin, heavy polypeptide 1, skeletal muscle, adult | 6.73 | 0.0404 | broad | motorprotein involved in spermiogenesis | (Wu et al. 2021) |
|  | *Actn2* | actinin alpha 2 | 6.40 | 0.0380 | broad | motorprotein involved in spermiogenesis | (Wu et al. 2021) |
|  | *Sln* | sarcolipin | 6.01 | 0.0307 | broad | no information on sperm effects |  |
|  | *Tnnt3* | troponin T3, skeletal, fast | 5.86 | 0.0050 | broad | motorprotein involved in spermiogenesis | (Wu et al. 2021) |
|  | *Mylk4* | myosin light chain kinase family, member 4 | 5.85 | 0.0138 | broad | motorprotein involved in spermiogenesis | (Wu et al. 2021) |

<sup>#1</sup> selected from the top 17 genes in the whole list (see suppl. Table1b)
<sup>#2</sup> selected from the top 27 genes in the whole list (see suppl. Table1d)
<sup>#3</sup> represent the top 10 genes from the list (see suppl. Table1f)

## Discussion

Based on comparative genomic and functional analysis, we show here that a new intra-intronic promotor has arisen in the mouse lineage 2-6 million years ago and has led to the evolution of a functionally new transcript within an otherwise highly conserved gene. The transcript is specific to the testis and knockout combined with transcriptome analysis shows that it is functionally involved in sperm maturation. Interestingly, it got also secondarily lost in some mouse lineages, apparently due to the acquisition of a disabling mutation in the promotor region.

The emergence of evolutionary novelties out of regulatory changes is by now well documented in many species (see (Carroll 2008, He et al. 2021, Mattioli et al. 2020, Osada et al. 2017, Romero et al. 2012, Signor and Nuzhdin 2018, Tautz 2000, Wray 2007) for a subset of relevant papers and reviews). In fact, it is so abundant that it constitutes often the first measurable differences in population and species divergence, including diverging mouse populations (Bryk et al. 2013), which raises the question whether much of is initially neutrally evolving or could be functional (Fay and Wittkopp 2008, Hill et al. 2021, Hodgins-Davis et al. 2019, Staubach et al. 2010). Given the evolutionary volatility of the Map2k7-α1 transcript, with its fast secondary loss after its initial emergence, one would normally have considered it to be mostly neutral and therefore subject to random fixation or loss. However, our data show that it has a clear functional role in spermatogenesis. The high evolutionary dynamics of this transcript is therefore more likely explained by the general effects of sexual selection that would be particularly effective in the germline and the gonads (Kleene 2005).

### Possible function of the Map2k7-α1 transcript

There are several possibilities of how the Map2k7-α1 transcript could function. The first is that it leads to the translation of a truncated protein that codes only for *Map2k7* kinase, but does not bind specifically to JNK. It could therefore phosphorylate other signaling proteins, but in an unspecific manner. This would likely be detrimental, rather than advantageous for the cells. Also, since we do not find GO terms that relate to signaling processes in the transcriptome analysis of knockout mice, we assume that the truncated protein is not expressed, or at least not funtional.

The second possibility is that Map2k7-α1 acts as a non-coding RNA. There are multiple ways of how non-coding RNAs can regulate other genes or gene complexes (Gil and Ulitsky 2020, Statello et al. 2021) and some have been implicated in male infertility (Joshi and Rajender 2020). In a previous study, we identified a testis specific new transcript that has also emerged via a new promotor acquisition and for which we could infer that it acts as lncRNA in spermiogenesis (Heinen et al. 2009). However, given that most of the Map2k7-α1 RNA overlaps with the functional *Map2k7* transcripts, it would seem unlikely that it could have assumed such a function as non-coding RNA as a whole, since most of its RNA is actually potentially coding. Only the new exon that emerged out of intronic sequences might have such a function.

On the other hand, the first AUG in this new transcript is embedded in an optimal Kozak-consensus-sequence (Kozak 1987) and one would therefore expect that it leads to the translation of a 50aa ORF. The resulting peptide does not match with any other protein or domain in the data bases, since it is actually produced out of a previously non-coding intron sequence.

While it has long been thought that proteins that emerge out of such more or less random sequences would not be functional, it has by now become clear that the *de novo* evolution of proteins is well possible (Tautz and Domazet-Loso 2011, Van Oss and Carvunis 2019). In fact, we have recently described such a case of a very recent *de novo* emergence of a protein that regulates pregnancy cycles in mice (Xie et al. 2019). In that case, we could proof a direct function of the protein in a knockout mouse which carried only a frameshift mutation in the protein. In the current study, we have deleted the whole transcript, implying that it is not fully proven that it is really the translated peptide that conveys the function. But studies with random peptide sequences in *E. coli* and in plants have shown that a substantial fraction of them can have a direct positive effect on their hosts (Bao et al. 2017, Neme et al. 2017). It seems therefore possible that the new peptide that is encoded by the Map2k7-α1 transcript is indeed a *de novo* protein with a function. Hence, this would be a case where a pre-existing potentially functional sequence was “waiting” for a promotor emergence to allow it to become functional.

The ORF is actually already present in the outgroup species, but would not expected to be translated in these species. The fact that it got secondarily lost in the *M. m. musculus* subspecies is in line with the observation of fast gain and loss cycles of *de novo* evolved transcripts and proteins (Neme and Tautz 2014, Palmieri et al. 2014). But our functional data show that despite of this evolutionary instability, the new transcript (and/or peptide) can still be functional.

## Acknowledgements

We thank Heike Harre for help with the sperm counting and phenotyping, Cornelia Burghardt for RNA extraction and the mouse team lead by Christine Pfeifle for expert support. This work was funded by institutional funds of the MPG to DT.

## Author contributions

T.H.: conceptualization, experimental work, data analysis, paper writing, C.X.: data analysis, M.K.: data analysis, D.S.: experimental work, data analysis, S.K.: experimental work, data analysis, D.T.: conceptualization, data analysis, paper writing. Part of this work was done in the framework of the PhD thesis of the first author (Heinen 2008).

## Supplementary Material

### Supplementary file S1

Exon specific Northen-Blotting and qPCR of *Map2k7*

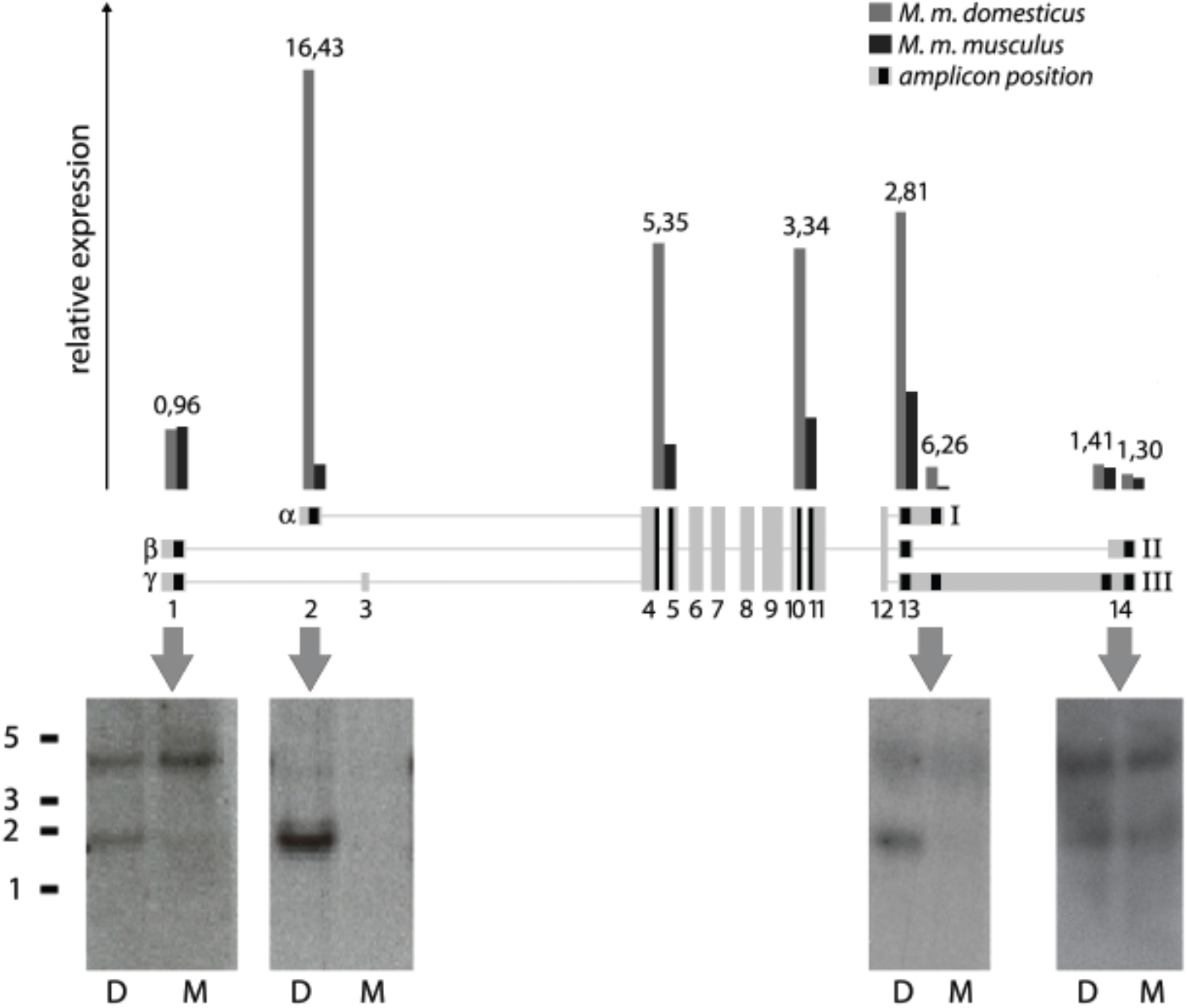

Exon specific *Map2k7* Northern blots and qRT-PCRs comparing *M. m. musculus* and *M. m. domesticus*. Testis RNA from *M. m. domesticus* (D) and *M. m. musculus* (M) was hybridized with different probes against certain parts of *Map2k7*. Probe positions of the respective blots are indicated with grey arrows. The size standard on the left displays kb. Relative expression of different qRT-PCR amplicons (about 100 bp in size) is shown at the top. Two amplicons span neighboring exons (4-5; 10-11). Positions of the different amplicons are indicated by black areas in the exon map. The respective expression values are displayed as bars above. The numbers on top of the bars represent the ratio of the domesticus to the musculus value. The results lead to the assumption of a strong prevalence of Map2k7-α1 in *M. m. domesticus* which is missing in *M. m. musculus*.

### Supplementary file S2

Targeting strategy for Map2k7-α1 knock out:

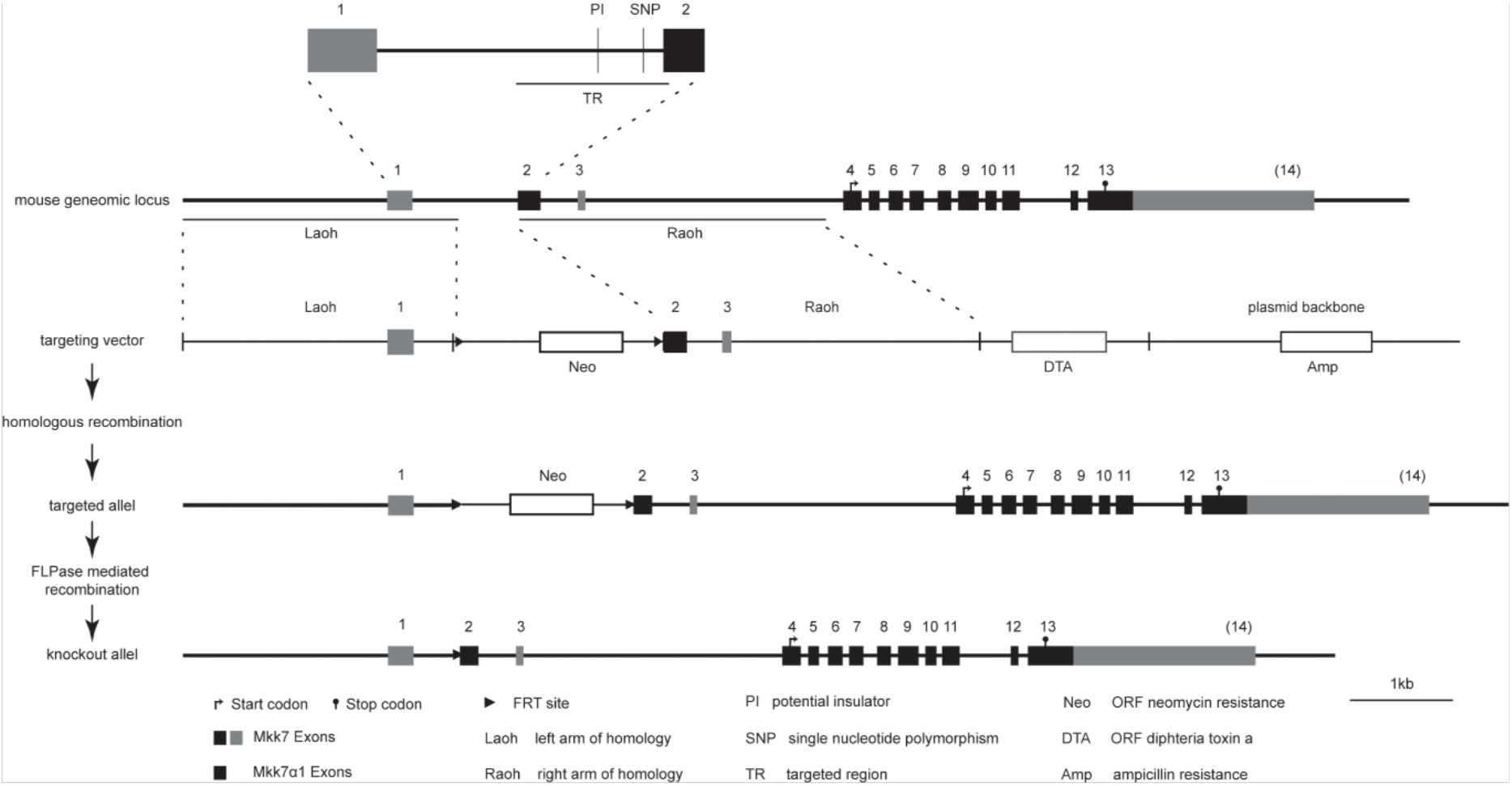

### Wildtype sequence

chr8:4,239,310- 4,240,429 in GRCm38/mm10

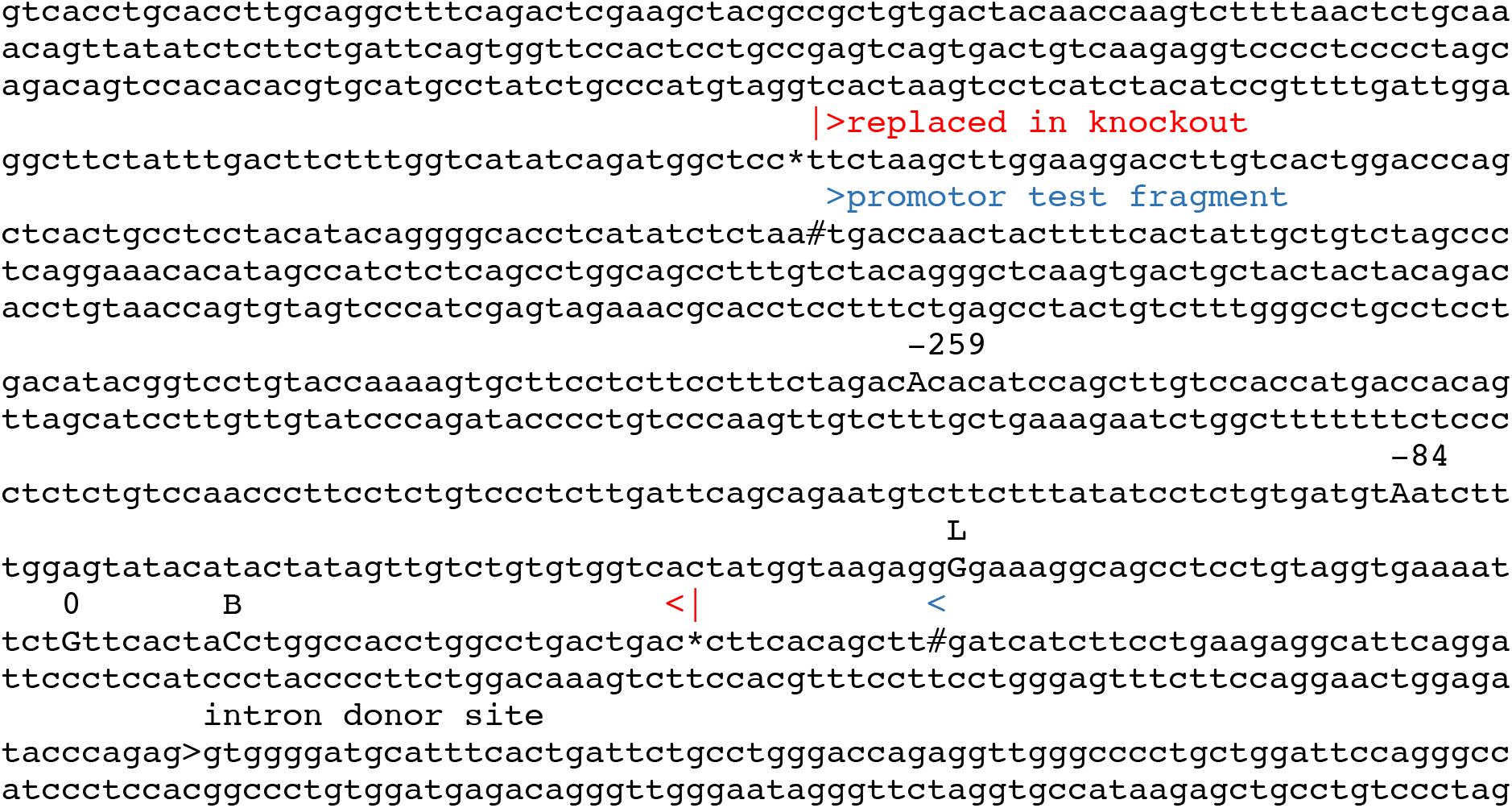

**Legend**: “0” marks the position that is used as reference for the transcription start site in Figure 4, “B” marks the start of the bulk of the transcripts detected in the RNA-Seq experiments, “L” marks the start of the longest annotated transcript in GRCm38/mm10. The region replaced in the knockout construct is indicated by * and red labels, the region used for promotor analysis is indicated by # and blue labels.

### Inserted NEO vector

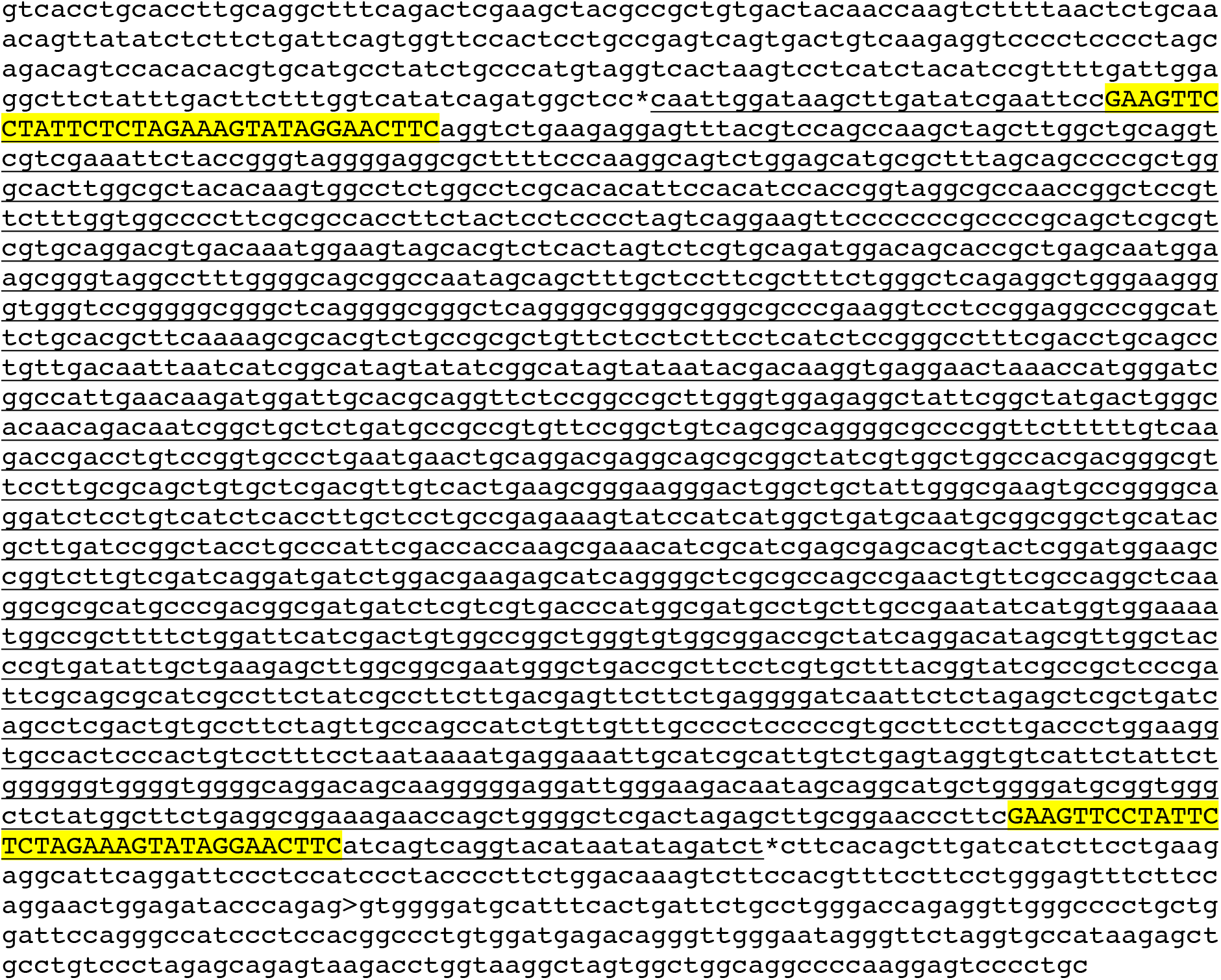

### Sequence after FRT recombination

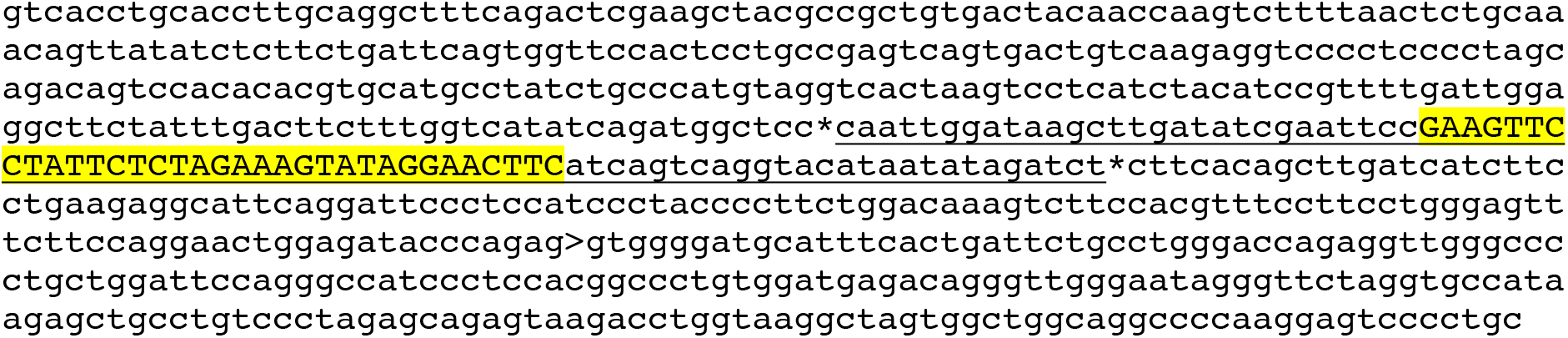

### vector derived sequences underlined

FRT sites in yellow

